# Identification of stable reference genes in *Edwardsiella ictaluri* for accurate gene expression analysis

**DOI:** 10.1101/2024.08.16.608217

**Authors:** Jingjun Lu, Hossam Abdelhamed, Nawar Al-Janabi, Nour Eissa, Mark Lawrence, Attila Karsi

## Abstract

*Edwardsiella ictaluri* is a Gram-negative bacterium causing enteric septicemia of catfish (ESC), leading to significant economic losses in the catfish farming industry. RT-PCR analysis is a powerful technique for quantifying gene expression, but normalization of expression data is critical to control experimental errors. Using stable reference genes, also known as housekeeping genes, is a common strategy for normalization, yet reference gene selection often lacks proper validation. In this work, our goal was to determine the most stable reference genes in *E. ictaluri* during catfish serum exposure and various growth phases. To this goal, we evaluated the expression of 27 classical reference genes (*16SrRNA*, *abcZ*, *adk*, *arc*, *aroE*, *aspA*, *atpA*, *cyaA*, *dnaG*, *fumC*, *g6pd*, *gdhA*, *glnA*, *gltA*, *glyA*, *grpE*, *gyrB*, *mdh*, *mutS*, *pgi*, *pgm*, *pntA*, *recA*, *recP*, *rpoS*, *tkt*, *and tpi*) using five analytical programs (GeNorm, BestKeeper, NormFinder, Comparative ΔCT, and Comprehensive Ranking). Results showed that *aspA*, *atpA*, *dnaG*, *glyA*, *gyrB*, *mutS*, *recP*, *rpoS*, *tkt*, and *tpi* were the most stable reference genes during serum exposure, whereas *fumC*, *g6pd*, *gdhA*, *glnA*, and *mdh* were the least stable. During various growth phases, *aspA*, *g6pd*, *glyA*, *gyrB*, *mdh*, *mutS*, *pgm*, *recA*, *recP*, *and tkt* were the most stable, while 16S rRNA, *atpA*, *grpE*, *and tpi* were the least stable. At least four analysis methods confirmed the stability of *aspA*, *glyA*, *gyrB*, *mutS*, *recP*, and *tkt* during serum exposure and different growth stages. However, no consensus was found among the programs for unstable reference genes under both conditions.

## Introduction

*Edwardsiella ictaluri* is a Gram-negative, rod-shaped, facultative anaerobic bacterium and the causative agent of enteric septicemia of catfish (ESC) (Hawke et al., 1981). ESC is a major disease that negatively impacts the catfish farming industry.

Real-time reverse transcription PCR (RT-PCR) has become an essential technique for quantifying gene expression in biological samples (Gibson et al., 1996; Purcell et al., 2011). RT-PCR offers high sensitivity and specificity, allowing for accurate expression profiling of selected targets in a short time (de Jonge et al., 2007; Eissa et al., 2017). This technology has facilitated the development of various assays for detecting and quantifying viral, parasitic, and bacterial fish pathogens (Bain et al., 2010; Griffin et al., 2009; Soto et al., 2010). In research, RT-PCR is used for measuring gene expression, estimating gene copy numbers, discovering new genes, and determining pathogen loads (Chuaqui et al., 2002; Giulietti et al., 2001; Higuchi et al., 1992).

Normalization of RT-PCR data using reference genes is crucial for accurate gene expression analysis and error control among samples (Suzuki et al., 2000; Tanic et al., 2007). An ideal reference gene should exhibit constant expression with minimal variation under different experimental conditions (Thellin et al., 1999). Using an unstable or suboptimal reference gene can lead to biased and ambiguous interpretations and conclusions (Dheda et al., 2005; Vandesompele et al., 2002). Studies have shown that classical reference genes demonstrate inconsistent expression across different experimental conditions (Tunbridge et al., 2011; Vandecasteele et al., 2001). Thus, using more than one reference gene is recommended to normalize RT-PCR data accurately (Andersen et al., 2004; Derveaux et al., 2010; Vandesompele et al., 2002).

Although many reference genes have been used in gene expression studies, no universal endogenous normalizer genes have been identified. Interestingly, RT-PCR-based gene expression studies in *E. ictaluri* are lacking, and to our best knowledge, there is no information on previously used reference genes or validation studies in this bacterium. Therefore, in this study, we aimed to identify stably expressed reference genes in *E. ictaluri* by assessing the expression of 27 classical reference genes under serum exposure and during different growth phases.

## Materials and methods

### Bacterial strains and culture conditions

*Edwardsiella ictaluri* strain 93-146 was used in this study. It was cultured in brain heart infusion (BHI) broth or on agar plates (Difco, Sparks, MD) and incubated at 30°C with shaking at 200 rpm. Colistin was added to the growth media at 12.5 μg/ml when needed.

### Reference genes, primer design, and PCR conditions

A total of 27 reference genes were selected from the literature based on the evolutionary distance between *E. ictaluri* and other bacteria (Maiden et al., 1998; Urwin and Maiden, 2003; Victor et al., 2007; Warsen et al., 2004). Table 1 contains a list of genes and their accession numbers.

**Table 1.** Reference genes in *E. ictaluri* being studied.

| Gene Symbol | ORF # | Product |
| --- | --- | --- |
| 16S rRNA | NT01EI_R00010 |  |
| *abcZ* | NT01EI_2798 | ABC transporter ATP-binding protein |
| *adk* | NT01EI_1123 | Adenylate kinase |
| *arcC* | NT01EI_3529 | Carbamate kinase |
| *aroE* | NT01EI_3555 | Dehydroshikimate reductase |
| *aspA* | NT01EI_0377 | Aspartase |
| *atpA* | NT01EI_3910 | ATP synthase alpha subunit |
| *cyaA* | NT01EI_0111 | adenylate cyclase |
| *dnaG* | NT01EI_0525 | DNA primase |
| *fumC* | NT01EI_2078 | Fumarate hydratase class II |
| *g6pd* | NT01EI_1607 | Glucose-6-phosphate dehydrogenase |
| *gdhA* | NT01EI_3732 | NADP-specific glutamate dehydrogenase |
| *glnA* | NT01EI_3863 | Glutamine synthetase |
| *gltA* | NT01EI_2875 | Citrate synthase |
| *glyA* | NT01EI_3190 | Serine hydroxy methyltransferase |
| *grpE* | NT01EI_3062 | Heat shock protein GrpE |
| *gyrB* | NT01EI_0004 | DNA gyrase subunit B |
| *mdh* | NT01EI_0446 | Malate dehydrogenase |
| *mutS* | NT01EI_3249 | Methyl-directed mismatch repair protein MutS |
| *pgi* | NT01EI_0210 | Glucose-6-phosphate isomerase |
| *pgm* | NT01EI_2893 | Phosphoglucomutase |
| *pntA* | NT01EI_1960 | Transhydrogenase alpha subunit |
| *recA* | NT01EI_3241 | RecA protein |
| *recP* | NT01EI_3370 | Transketolase |
| *rpoS* | NT01EI_3250 | RNA polymerase, sigma factor RpoS |
| *tkt* | NT01EI_3370 | Transketolase |
| *tpi* | NT01EI_3785 | Triosephosphate isomerase |

Forward and reverse primers were designed for each gene (Table 2) using Primer 3 software to amplify a product size of 100 to 150 nucleotides. Primer specificity was checked by PCR (AB 2720, Applied Biosystems) using *E. ictaluri* strain 93-146 genomic DNA. PCR conditions were: 94°C for 2 min; 35 cycles of 94°C for 30 s, 56°C for 30 s, and 72°C for 30 s; final extension at 72°C for 8 min, followed by a hold at 12°C. Amplified products were verified by agarose gel electrophoresis.

**Table 2.**
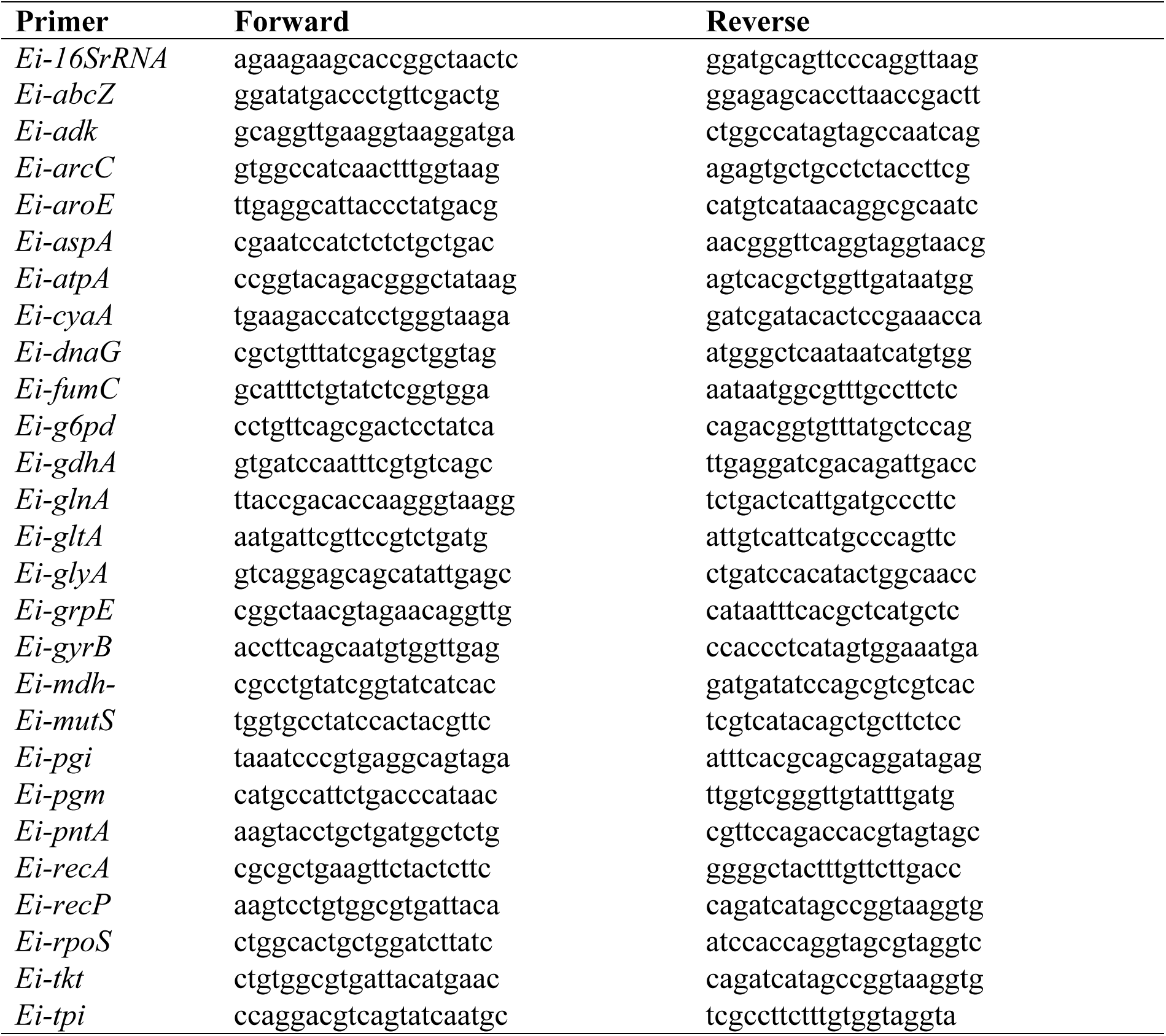
Primer pairs used in this study.

### Exposure of *E. ictaluri* to catfish serum

Serum was collected from 1-2 kg specific pathogen-free (SPF) channel catfish obtained from the fish hatchery at the College of Veterinary Medicine, Mississippi State University. Briefly, catfish were anesthetized in water containing 200 mg/liter of tricaine methanesulfonate (Argent Laboratories), and blood was collected from the caudal vein at 1% of the body weight. Then, blood samples were allowed to clot at room temperature for 30 min and placed on ice for an additional 30 min. Normal serum (NS) was collected by centrifugation with a Sorvall Legend RT centrifuge (Thermo Electron Corp., Asheville, NC) at 3000 rpm for 10 min 4°C, pooled, and stored at -80°C in aliquots of 1 ml. Heat-inactivated serum (HIS) was obtained by incubating NS at 65°C for 45 min. Before use, serum was allowed to thaw on ice and then placed in a 30°C water bath for 15 min for a uniform temperature.

For serum exposure, four different colonies of *E. ictaluri* were inoculated separately in 2 ml BHI broth and grown at 30°C in a shaking incubator for 16 h. The optical densities (OD_600_) were measured using a Spectronic GENESYS 20 spectrophotometer (Thermo Electron, Waltman, MA) and adjusted to 1.0. Fifteen ml fresh media were inoculated with the overnight culture at 1:100 dilution and grown at 30°C until OD_600_ reached 1.0. Ten ml cultures were harvested by centrifugation at 4000 rpm for 10 min at 30°C and washed twice with standard cell wash buffer (10 mM Tris pH 8.0; 5 mM magnesium acetate). Before the third wash, dissolved bacteria in each of the four tubes were divided equally into another tube, yielding two sets of four pellets. 1.25 ml NS and HIS were added into sets one and two, respectively. After dissolving pellets in serum, tubes were incubated at 30°C for 2 h. During the incubation, tubes were inverted ten times every 30 min to mix bacteria and serum. Then, bacteria and serum mixtures were transferred into 15 ml centrifuge tubes containing 3 ml of RNAprotect Bacteria Reagent (Qiagen). After vortexing for 5 s at high speed, the mixture was incubated at room temperature for 10 min and aliquoted into four tubes, which were stored at -80°C until total RNA isolation.

### *E. ictaluri* growth kinetics

Small and large broth cultures from 4 different *E. ictaluri* colonies were prepared as described above, and the cultures’ OD_600_ values were determined at 3, 6, 12, 18, 24, and 30 h. Five ml culture was removed at 6, 9, 12, 24, 36, and 48 h and harvested by centrifugation at 4000 rpm for 10 min at 30°C. The pellets were resuspended in 3 ml RNAprotect Bacteria Reagent (Qiagen), aliquoted, and stored at -80°C until total RNA isolation.

### Total RNA isolation and cDNA synthesis

Total RNA was isolated from *E. ictaluri* following serum exposure and at different growth stages using RNeasy Protect Bacteria Mini Kit (Qiagen) according to the manufacturer’s protocol. Total RNA quality and quantity from each sample were measured using a NanoDrop 1000 (Thermo Scientific) and an Agilent Bioanalyzer (Agilent Technologies). First-strand cDNA was synthesized from 1 µg of total RNA with QuantiTect Reverse Transcription Kit (Qiagen). Double-strand cDNA was synthesized using the SuperScript Double-Stranded cDNA Synthesis Kit (ThermoFisher).

### Expression analysis of reference genes

After verification of primer specificity and amplicon size, expression of reference genes was determined from 10-fold diluted cDNAs using a Stratagene Mx3005P qPCR system (Agilent Technologies) with QuantiTect SYBR Green RT-PCR Kit (Qiagen). Reactions were conducted in a total volume of 20 μl in a 96-well optical reaction plate (Stratagene). RT-PCR conditions were as follows: 95°C for 10 min; 40 cycles of 95°C for 15 s, 55°C for 30 s, 72°C for 30 s. A dissociation curve analysis was performed after 40 cycles using the following conditions: 95°C for 1 min, 55°C for 30 s, and 95°C for 30s.

### Stability analysis of reference genes

Raw CT values from qPCR were analyzed using five programs (geNorm, BestKeeper, NormFinder, Comparative ΔCT, and comprehensive gene ranking). geNorm calculates the average expression stability value (M value)based on intragroup differences and mean pairwise variation (Vandesompele et al., 2002). BestKeeper evaluates stability based on the standard deviation (SD) and coefficient of variation (CV) of raw CT values (Pfaffl et al., 2004).

NormFinder estimates intragroup and intergroup variation and combines the two values into a stability value ρ (the lowest ρ is the most stable) (Andersen et al., 2004). Comparative ΔCT compares the relative expression of “pairs of genes” within each sample (Silver et al., 2006). Finally, the comprehensive ranking tool integrates results from all methods by assigning weights to each reference gene and calculating the geometric mean for the final ranking (Xie et al., 2011).

## RESULTS

### Edwardsiella ictaluri growth kinetics

The growth curve of *E. ictaluri* in BHI broth was determined by measuring the OD_600_ at 3, 6, 12, 18, 24, and 30 h. As shown in Fig 1, *E. ictaluri* reached the early log phase at approximately 6 h, and entered the stationary phase at approximately 24 h. Based on this growth curve, bacterial samples were collected at early log phase (6 h), middle log phase (9 h), late log phase (12 h), early stationary phase (24 h), middle stationary phase (36 h), and late stationary phase (48 h).

**Fig 1.**
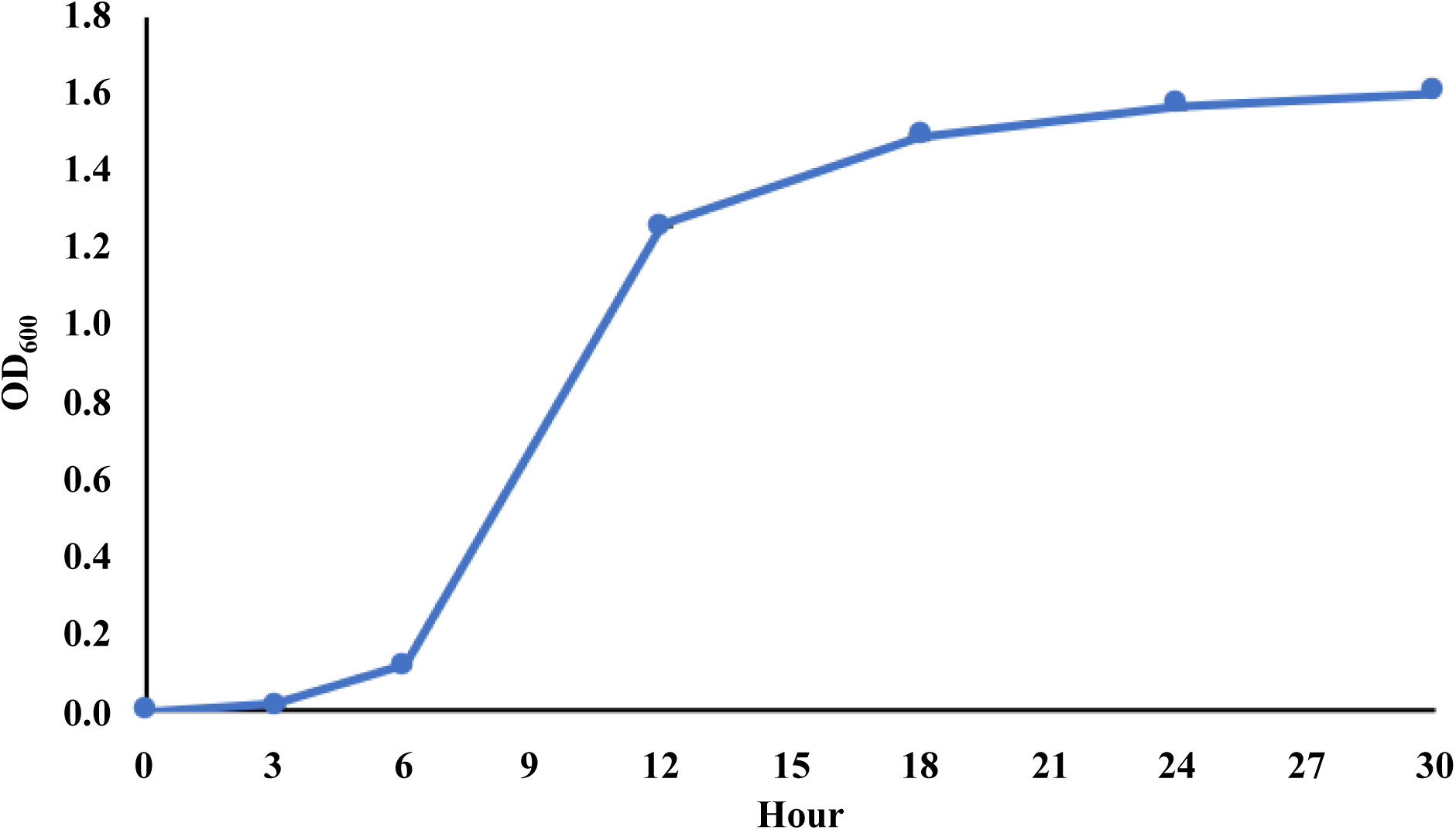
Growth kinetics of *E. ictaluri* in BHI broth.

### Expression of reference genes during serum exposure and growth

The expression levels of the 27 reference genes were evaluated using CT and descriptive statistics (min, max, median, mean, and SD). In the serum exposure experiment (Table 3), the mean CT values for reference genes ranged between 24.9 and 35.05. *fumC* had the highest median CT value (37.11), which indicated a relatively low expression level, while *dnaG* had the lowest median CT value (CT = 27.06), which indicated a relatively high expression level. In the growth phase experiment (Table 4), the mean CT values for the tested genes ranged between 9.74 and 33.5. *fumC* had the highest median CT value (32.65), which indicated a relatively low expression level, while *16s RNA* had the lowest median CT value (CT = 9.79), which indicated relatively high expression.

**Table 3.** Descriptive statistics of 27 reference genes in *E. ictaluri* exposed to serum.

| Genes | Minimum | Maximum | Median | Mean | Std. Deviation |
| --- | --- | --- | --- | --- | --- |
| 16S rRNA | 23.08 | 32.29 | 31.21 | 29.09 | 3.682 |
| *grpE* | 25.9 | 34.54 | 33.88 | 31.51 | 3.441 |
| *aroE* | 25.03 | 33.84 | 32.78 | 30.68 | 3.442 |
| *abcZ* | 25.6 | 34.44 | 33.63 | 31.4 | 3.525 |
| *adk* | 26.56 | 35.34 | 33.58 | 31.97 | 3.385 |
| *tpi* | 24.89 | 33.45 | 32.42 | 30.4 | 3.405 |
| *arcC* | 19.85 | 28.02 | 26.67 | 24.99 | 3.325 |
| *aspA* | 20.24 | 29.07 | 27.76 | 25.72 | 3.496 |
| *cyaA* | 24.7 | 34.19 | 32.82 | 30.46 | 3.777 |
| *atpA* | 21.04 | 29.42 | 28.63 | 26.45 | 3.353 |
| *fumC* | 28.15 | 40.00 | 37.11 | 35.05 | 4.909 |
| *dnaG* | 19.77 | 27.89 | 27.06 | 24.9 | 3.281 |
| *gdhA* | 24.97 | 40.00 | 31.13 | 30.99 | 4.843 |
| *g6pd* | 26.62 | 40.00 | 32.31 | 32.06 | 4.399 |
| *gltA* | 29.43 | 40.00 | 34.99 | 34.59 | 3.337 |
| *glnA* | 26.45 | 40.00 | 32.11 | 31.89 | 4.428 |
| *mdh* | 26.08 | 40 | 31.68 | 31.5 | 4.606 |
| *gyrB* | 25.43 | 31.97 | 30.81 | 29.1 | 2.966 |
| *mutS* | 25.21 | 33.99 | 33.09 | 31.22 | 3.36 |
| *glyA* | 23.91 | 32.65 | 31.53 | 29.35 | 3.507 |
| *pgm* | 28.82 | 35.26 | 34.26 | 32.29 | 3.098 |
| *pgi* | 27.31 | 33.59 | 32.62 | 30.76 | 2.984 |
| *recA* | 24.64 | 30.89 | 29.62 | 28 | 2.92 |
| *pntA* | 24.2 | 33.08 | 32.23 | 29.87 | 3.629 |
| *rpoS* | 24.73 | 33.27 | 32.19 | 30.06 | 3.521 |
| *recP* | 25.18 | 33.29 | 32.73 | 30.43 | 3.326 |
| *tkt* | 25.21 | 33.97 | 32.72 | 30.67 | 3.478 |

**Table 4.** Descriptive statistics of 27 reference genes in *E. ictaluri* growth.

| Genes | Minimum | Maximum | Median | Mean | Std. Deviation |
| --- | --- | --- | --- | --- | --- |
| *abcZ* | 26.16 | 35.80 | 30.60 | 30.70 | 2.43 |
| *adk* | 25.88 | 38.47 | 31.19 | 31.99 | 3.45 |
| *aspA* | 18.40 | 28.85 | 22.28 | 22.97 | 3.27 |
| *arcC* | 21.38 | 33.41 | 24.24 | 25.57 | 3.47 |
| *g6pd* | 24.24 | 34.83 | 27.86 | 28.20 | 2.98 |
| *gdh* | 25.76 | 36.38 | 28.20 | 29.73 | 3.47 |
| *dnaG* | 20.48 | 30.62 | 24.06 | 24.38 | 2.66 |
| *fumc* | 29.61 | 40.00 | 32.65 | 33.55 | 3.21 |
| *atpA* | 20.14 | 35.37 | 25.96 | 26.39 | 4.92 |
| *cyaA* | 29.16 | 37.30 | 32.86 | 32.49 | 2.29 |
| *tkt* | 21.92 | 31.66 | 26.68 | 26.66 | 3.33 |
| 16S rRNA | 8.57 | 11.75 | 9.79 | 9.74 | 0.70 |
| *recP* | 22.14 | 31.98 | 26.96 | 27.02 | 3.28 |
| *rpos* | 24.06 | 32.86 | 26.41 | 27.45 | 2.47 |
| *pntA* | 22.39 | 33.28 | 25.78 | 26.87 | 3.64 |
| *recA* | 22.69 | 30.70 | 25.73 | 26.17 | 2.74 |
| *pgi* | 22.49 | 34.94 | 27.02 | 27.34 | 3.96 |
| *pgm* | 24.74 | 35.11 | 29.00 | 29.17 | 3.34 |
| *gyrB* | 22.74 | 31.42 | 25.88 | 26.31 | 2.92 |
| *mdh* | 20.22 | 29.80 | 22.28 | 23.64 | 3.00 |
| *glyA* | 26.59 | 35.87 | 29.59 | 30.15 | 2.67 |
| *mutS* | 22.13 | 32.95 | 25.87 | 26.74 | 3.56 |
| *glna* | 21.24 | 34.43 | 25.45 | 26.45 | 4.48 |
| *gltA* | 23.46 | 34.64 | 26.66 | 28.07 | 3.20 |
| *aroE* | 26.94 | 36.26 | 30.42 | 30.62 | 2.66 |
| *grpE* | 22.61 | 34.89 | 28.36 | 27.45 | 3.68 |
| *tpi* | 22.24 | 34.69 | 28.27 | 28.16 | 4.25 |

### Stability of reference genes

The expression stability of 27 reference genes was evaluated under the tested conditions by geNorm, BestKeeper, NormFinder, comparative ΔCt, and comprehensive ranking system. Results are provided below for each analysis.

### GeNorm

The geNorm algorithm determines the stability of reference genes based on expression stability value (M). Gene with the lowest stability value is expressed most stably. This cut-off value of 1.5 has been adopted widely as a criterion for selecting reference genes (Ling and Salvaterra, 2011; Maroufi et al., 2010; Van Hiel et al., 2009). Under serum stress, all genes except *mdh* and *gdhA* had stability values below 1.5. The top 5 stable reference genes were *aspA* and *tkt*, *tpi*, *rpoS*, and *pntA* while *gltA*, *g6pd*, *glnA*, *mdh*, and *gdhA* were the least stable reference genes (Fig 2A). For the growth experiment, 19 reference genes were below the 1.5 threshold. The top 5 stable reference genes were *tkt* and *recP*, *mutS*, *pgm*, and *recA* while *adk*, *abcZ*,16S rRNA, *grpE*, and *tpi* were the least stable reference genes (Fig 2B).

**Fig 2.**
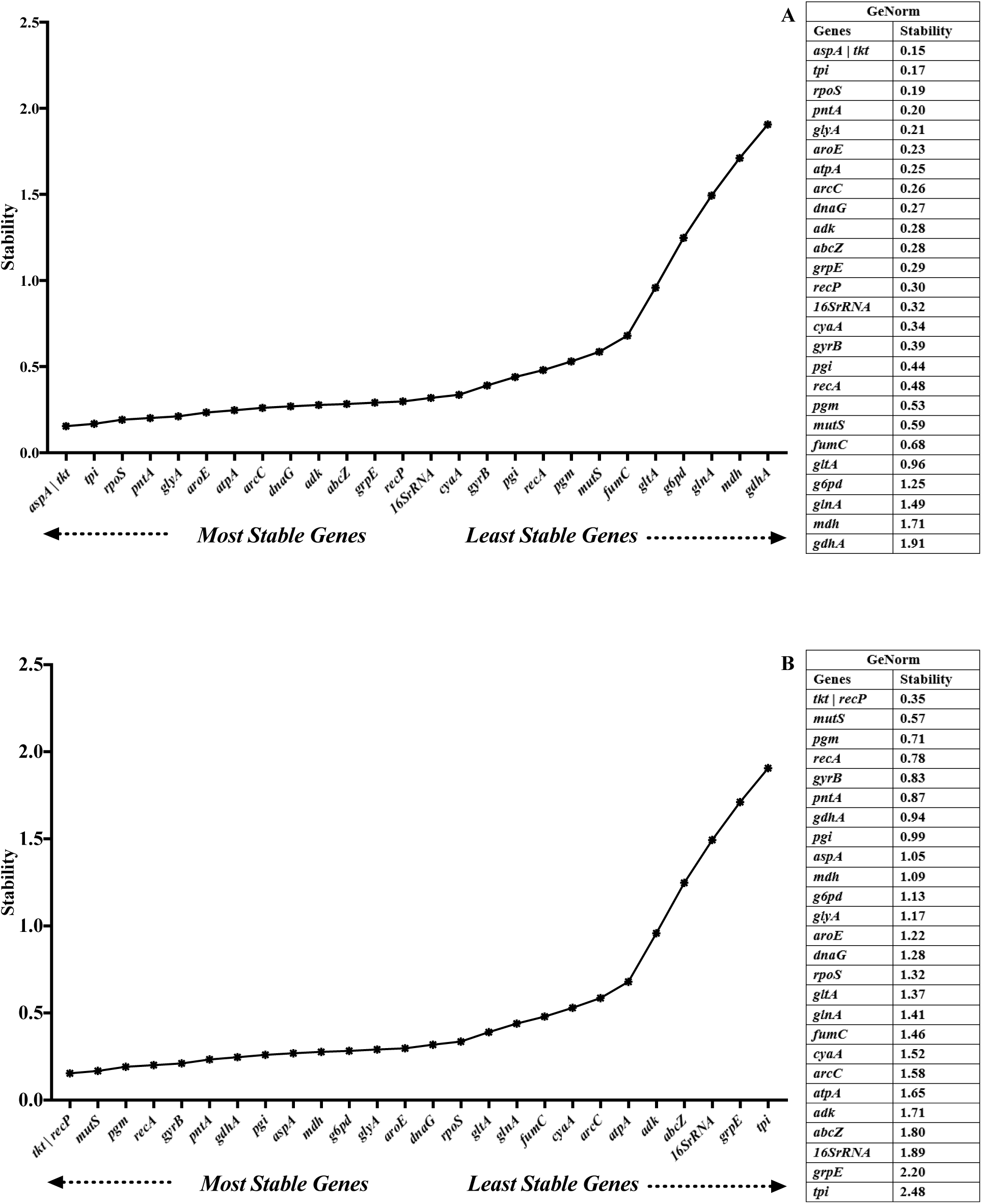
Expression stability of reference genes under serum exposure (A) and different growth phases (B) analyzed by GeoNorm. Lower M values indicate higher stability.

### BestKeeper

In the serum exposure experiment, all *E. ictaluri* reference genes lacked stability (SD > 1.00) (Fig 3A). In the growth phase experiment, all *E. ictaluri* reference genes, except *glnA* displayed good stability (SD < 1.00). The top 5 stable reference genes were 16S rRNA, *abcZ*, *adk*, *arcC*, and *cyaA* while *mutS*, *gyrB*, *pgm*, *pntA*, and *glna* were the least stable reference genes (Fig 3B).

**Fig 3.**
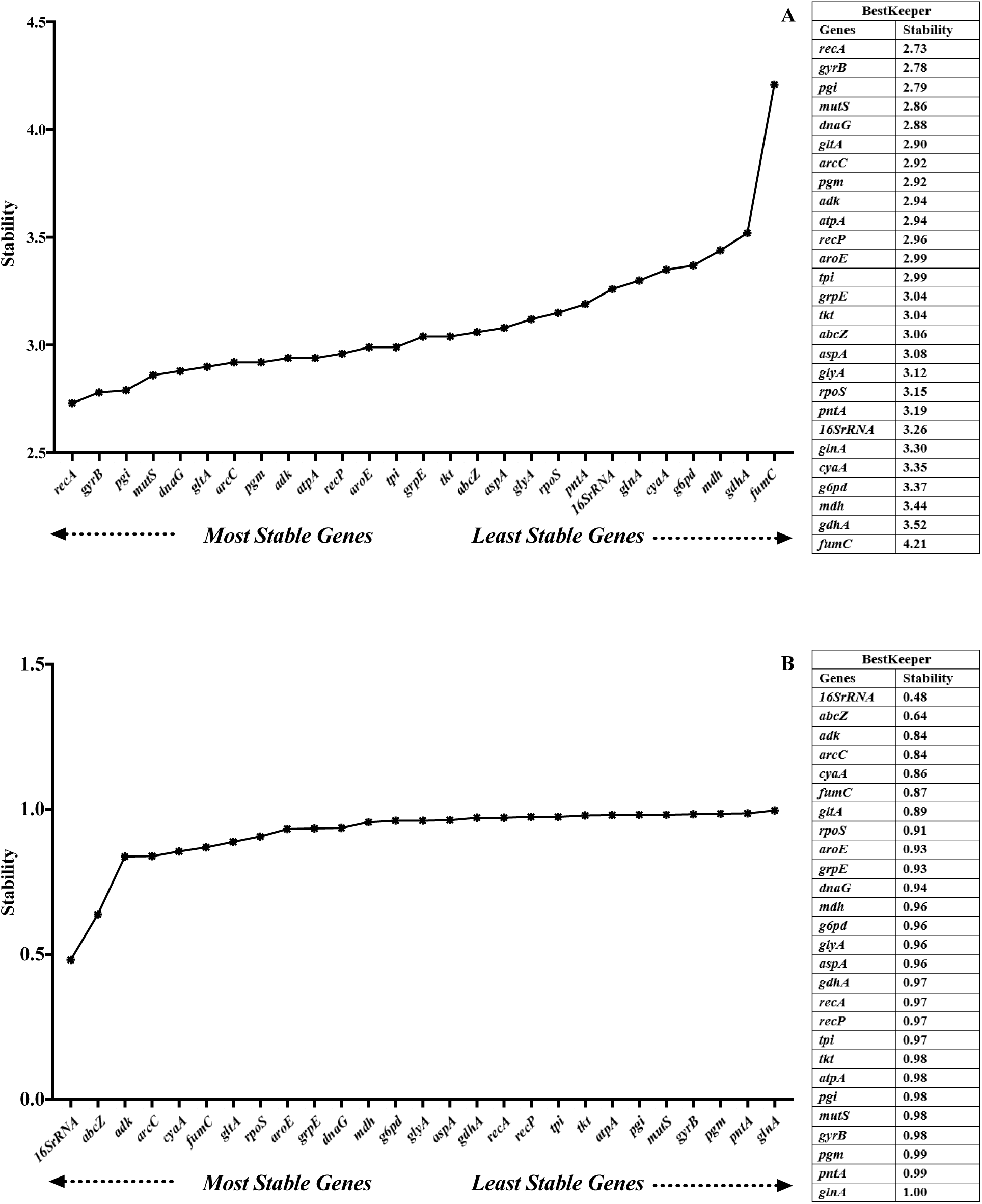
Expression stability of reference genes under serum exposure (A) and different growth phases (B) analyzed by BestKeeper gene.

### Normfinder

Lower average stability values indicate more stable and optimal expression. In serum exposure, the top 5 stable reference genes were *mutS*, *recP*, *tpi*, *arcC*, and *tkt*, while *fumC*, *gltA*, *g6pd*, *glnA*, *mdh*, and *gdhA* were the least stable (> 2) (Fig 4A). In growth phases, the top 5 stable reference genes were *gyrB*, *pgm*, *glyA*, *recA*, and *mdh,* whereas *atpA*, *abcZ*, 16S rRNA, *grpE*, and *tpi* were among the least stable genes (> 2) (Fig 4B).

**Fig 4.**
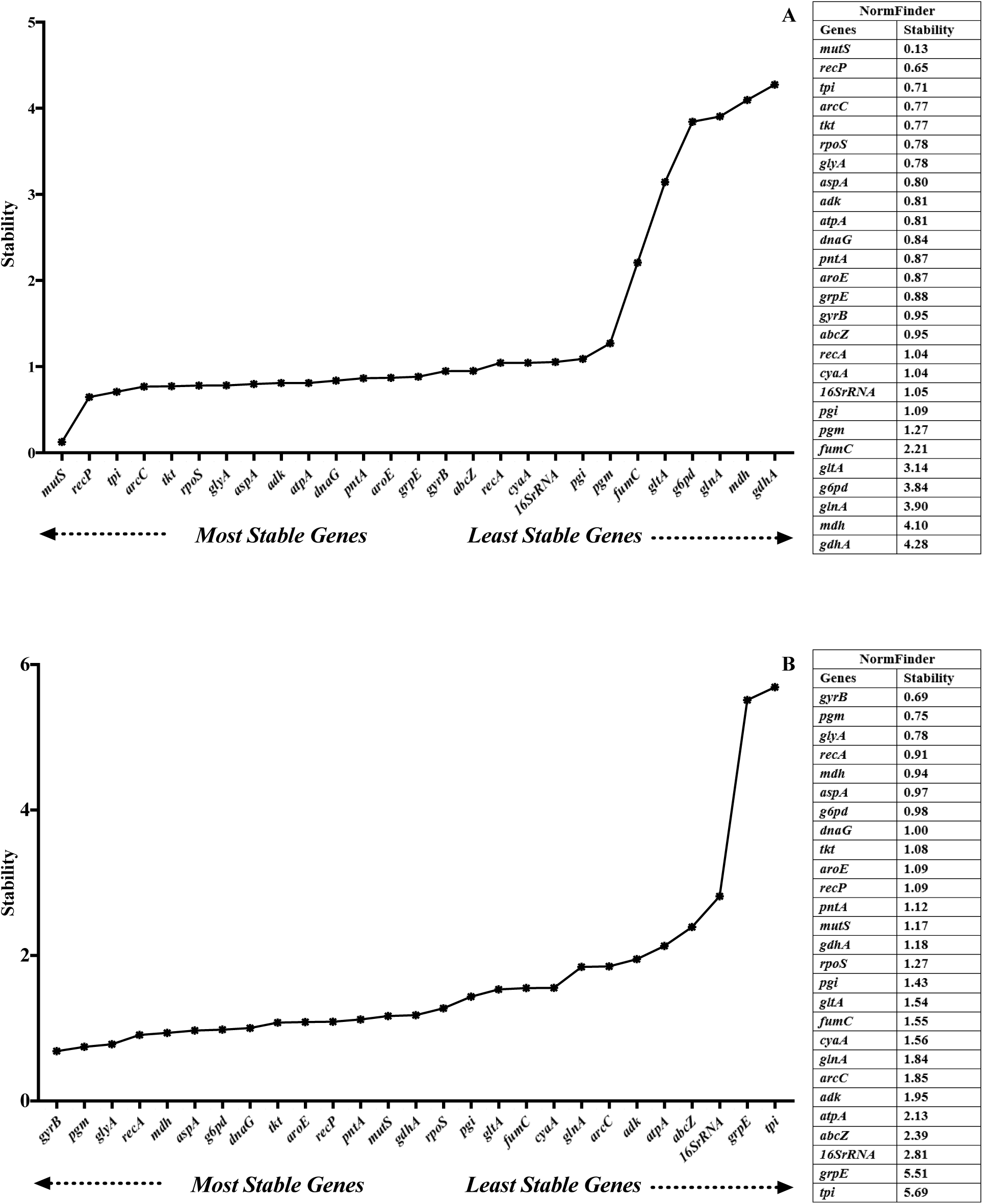
Expression stability of reference genes under serum exposure (A) and different growth phases (B) analyzed by NormFinder.

### Comparative **Δ**CT

The ΔCT method ranks all genes’ stability based on the repeatability of the gene expression differences among various samples (STDEV average). In serum exposure, the top 5 stable reference genes were *tpi*, *tkt*, *aspA*, *rpoS*, and *atpA*, while *fumC*, *gltA*, *g6pd*, *glnA*, *mdh*, and *gdhA* were among the least stable reference genes (> 2) (Fig 5A). In growth phases, the top 5 stable reference genes were *gyrB*, *pgm*, *recA*, *mdh*, and *tkt*, whereas 16S rRNA, *grpE*, and *tpi* showed the lowest expression stability (> 3) (Fig 5B).

**Fig 5.**
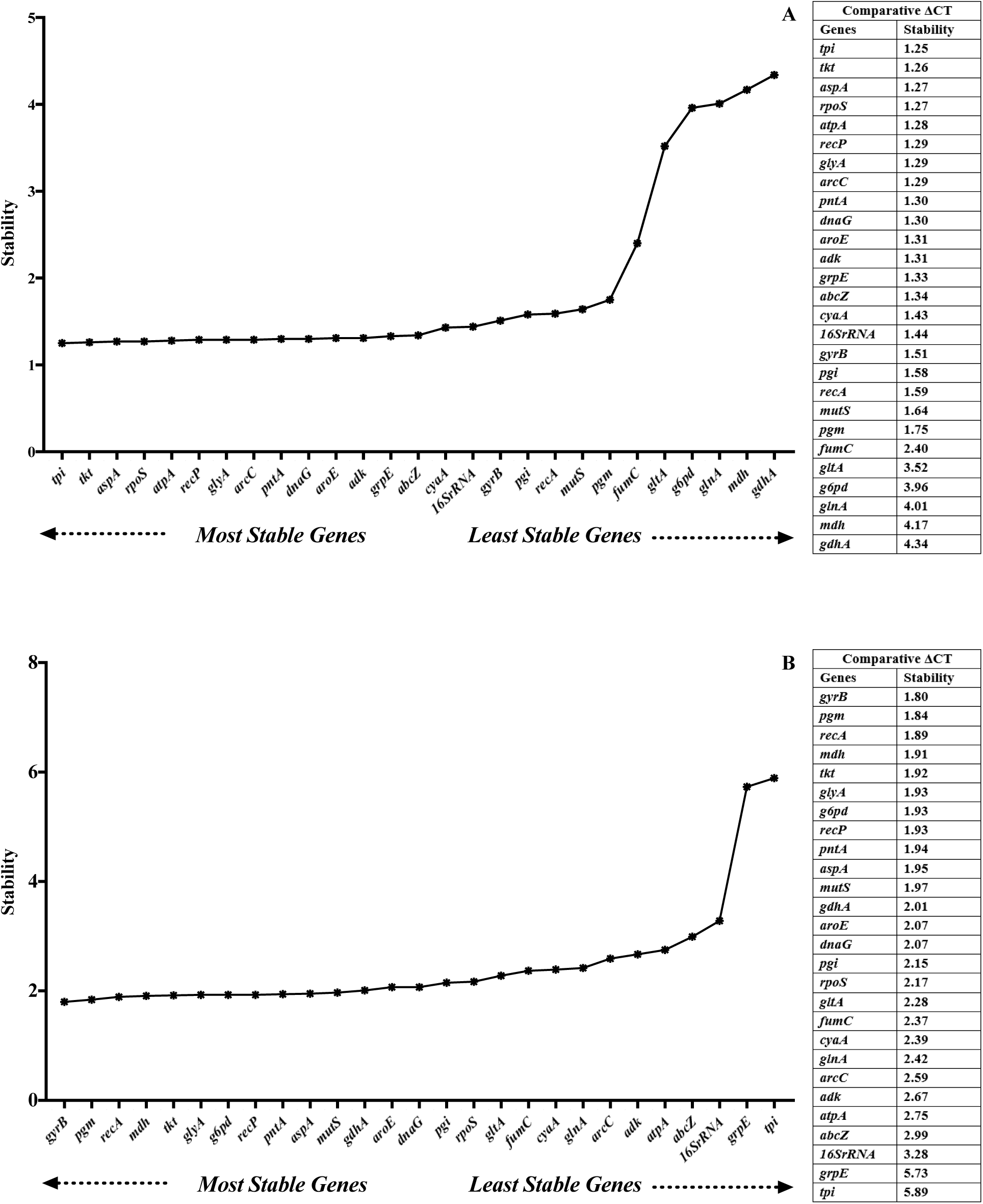
Expression stability of reference genes under serum exposure (A) and different growth phases (B) analyzed by Comparative ΔCT method.

### Comprehensive gene ranking

The results obtained from geNom, BestKeeper, NormFinder and comparative ΔCT were analyzed using the comprehensive ranking tool. In serum exposure, the top 5 stable reference genes were *tpi*, *tkt*, *aspA*, *mutS*, and *rpoS* while *fumC*, *gltA*, *g6pd*, *glnA*, *mdh*, and *gdhA* were among the least stable reference genes (Fig 6A). The top 5 stable reference genes in different growth phases were *gyrB*, *pgm*, *recA*, *tkt*, and *recP,* whereas 16S rRNA, *grpE*, and *tpi* showed the lowest expression stability (Fig 6B).

**Fig 6.**
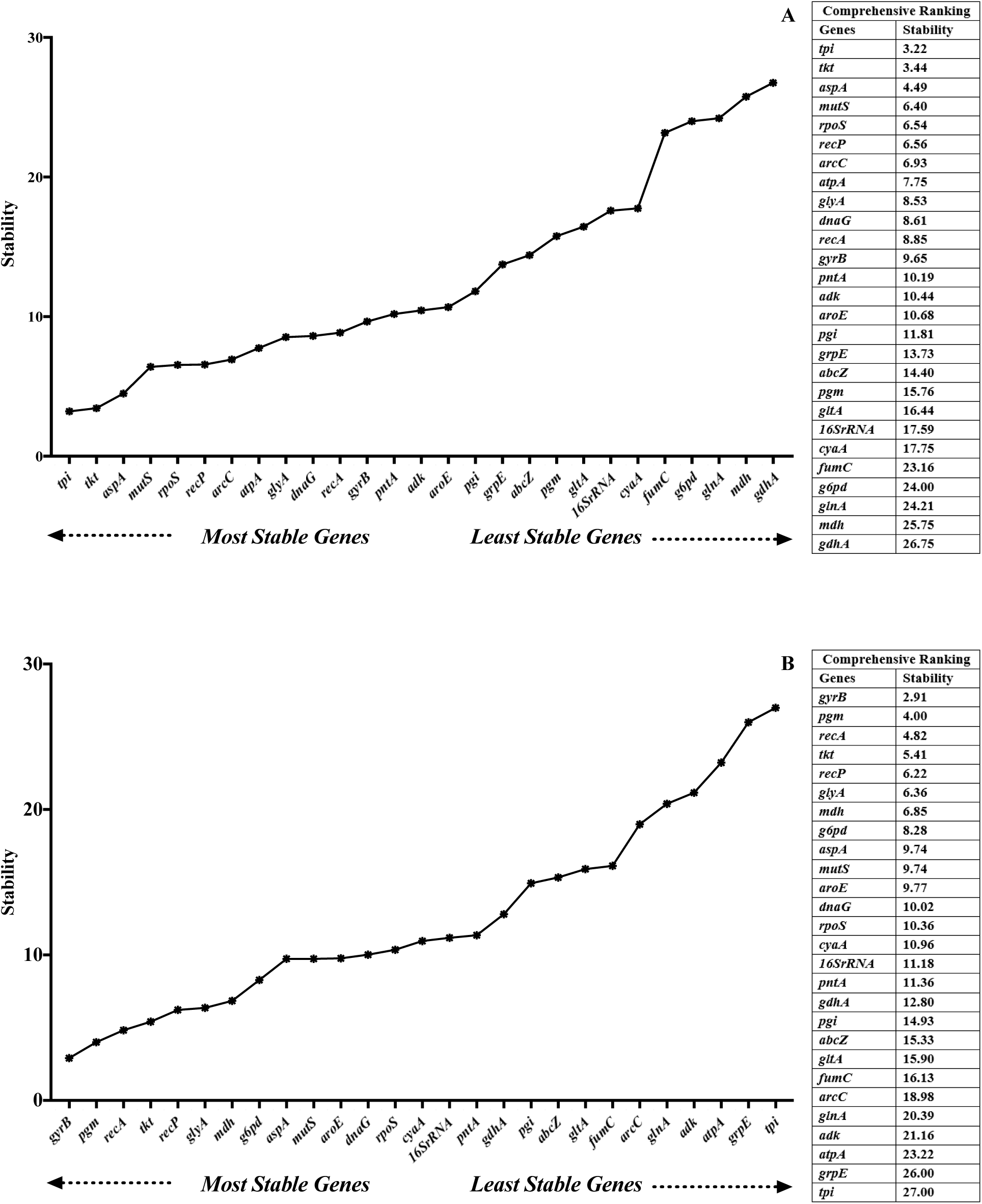
Comprehensive ranking of reference gene stability under serum exposure (A) and different growth phases (B).

### Most and least stable reference genes

During *E. ictaluri* serum exposure, computational programs consistently identified *aspA*, *atpA*, *dnaG*, *glyA*, *gyrB*, *mutS*, *recP*, *rpoS*, *tkt*, and *tpi* as the most stable reference genes (identified by at least 4 programs). Conversely, *fumC*, *g6pd*, *gdhA*, *glnA*, and *mdh* were the least stable reference genes (identified by at least 3 programs). All five programs indicated that *dnaG* was stable and *gdhA* and *mdh* were not stable in *E. ictaluri* during serum exposure.

During different *E. ictaluri* growth phases, computational programs consistently identified *aspA*, *g6pd*, *glyA*, *gyrB*, *mdh*, *mutS*, *pgm*, *recA*, *recP*, *and tkt* as the most stable reference genes (identified by at least 4 programs). Conversely, 16S rRNA, *atpA*, *grpE*, *and tpi* were the least stable reference genes (identified by at least 3 programs). All five programs indicated that *aspA*, *g6pd*, *gyrB*, *mdh*, *pgm*, and *recA* were stable, while all programs except BestKeeper indicated *grpE* and *tpi* were not stable during *E. ictaluri* growth.

## Discussion

Gene expression analysis is essential for investigating gene activity in living organisms. Stable reference genes are crucial for controlling errors and variations in input RNA in RT-PCR. Several studies have shown that expression of many commonly used reference genes is affected by growth phase, metabolic conditions, or experimental conditions (Dheda et al., 2004; Huggett et al., 2005; Schmittgen and Zakrajsek, 2000). Experimental conditions causing variation in expression of reference genes result in altered findings in target gene expression. Therefore, accurate quantification of target gene expression requires validation of reference gene stability (Huggett et al., 2005), which is often overlooked (Kozera and Rapacz, 2013). In this study, our goal was to determine stably expressed reference genes in *E. ictaluri* under serum exposure and various growth phases by assessing the expression of 27 classical reference genes.

The classical reference genes used in bacteria include 16S rRNA, *abcZ*, *adk*, *arc*, *aroE*, *aspA*, *atpA*, *cyaA*, *cysS*, *dnaG*, *fumC*, *g6pd*, *glcK*, *glnA*, *gltA*, *glyA*, *gmk*, *groEL*, *grpE*, *gyrA*, *gyrB*, *mdh*, *mutS*, *pgi*, *pgm*, *pntA*, *purB*, *recA*, *recP*, *rpoB*, *rpoD*, *rpoS*, *sghA*, *tkt*, *tpi* (Cusick et al., 2015; Florindo et al., 2012; Stenico et al., 2014; Vandecasteele et al., 2001; Zhongyang Sun et al., 2017). Serum stress and bacterial growth can affect gene expression in *E. ictaluri*. Thus, these conditions should provide a good assessment of the stability of reference genes. To this goal, expression of 27 classical reference genes was analyzed by five different programs because there is no consensus on which program is optimum for selecting reference genes (Cappelli et al., 2008; Kozera and Rapacz, 2013). Comparison of outcomes of different tools can yield significantly better candidates and lower the risk of artificial selection of co-regulated transcripts (Ayers et al., 2007).

When both serum exposure and growth phases of *E. ictaluri* were considered, at least four analysis methods indicated that *aspA*, *glyA*, *gyrB*, *mutS*, *recP*, and *tkt* reference genes were stable under both experimental conditions. However, the analysis methods did not reveal consensus unstable reference genes under either experimental condition.

We have not identified any publication reporting stability of *aspA* as a reference gene. However, when glucose, glycerol, and acetate were used as a carbon source in minimal media, expression of *Escherichia coli aspA* gene increased only in minimal media with glycerol and acetate during the exponential growth phase (Oh and Liao, 2000), which indicates that *aspA* gene expression could change under different experimental conditions. *glyA* was a suggested reference gene in medium with and without iron during the exponential and early stationary growth of *Actinobacillus pleuropneumoniae* (Nielsen and Boye, 2005). Stability of *gyrA* was verified in two different media at the mid-exponential growth phase for *Corynebacterium pseudotuberculosis*, but *gyrB* was excluded from the stability study due to low amplification linearity (Carvalho et al., 2014). *gyrB* was among the most stably expressed genes in *Staphylococcus aureus* under osmotic and acidic stress conditions (Sihto et al., 2014). It was also stably expressed in *Clavibacter michiganensis* under nutrient-rich and host-interaction conditions, as well as during a viable nonculturable state (Jiang et al., 2019). We have not identified any publication reporting the stability of *mutS* as a reference gene. However, in wild-type *E. coli* O157 : H7, the amount of MutS protein decreased about 26-fold in the stationary phase compared to the exponential phase (Li et al., 2003), which implies that *mutS* gene expression may change during bacterial growth. Information on the stability of *mutS* and *recP* as reference genes is not available in the literature. Finally, the expression of *tkt* changed at different temperature conditions in *Streptococcus agalactiae* (Florindo et al., 2012).

Analysis tools indicated that 16S RNA gene expression was more stable during serum exposure than *E. ictaluri* growth, but in general, it was not among the most stable reference genes. In contrast to our findings, 16S rRNA expression was the most stable reference gene in *Edwardsiella tarda* at different growth phases (Zhongyang Sun et al., 2017). 16S rRNA was also the most stable gene in *Shewanella psychrophila* at different hydrostatic pressures, but it was not an optimal reference gene under different temperatures and salinities (Liu et al., 2018). Similarly, 16S rRNA was the least reliable reference gene in *Bifidobacterium adolescentis* exposed to bile extract (Stenico et al., 2014). The usefulness of the 16S rRNA gene as a control is often doubted (Vetrovsky and Baldrian, 2013). These differences may exist due to sampling at different stages of bacterial growth because many housekeeping genes, as well as 16S rRNA, show increased expression before the mid-exponential growth phase, and 16S rRNA gene decreases significantly earlier than other reference genes at later stages of growth (Vandecasteele et al., 2001)

In this study, contradictory results were obtained regarding the stability of some reference genes between serum exposure and growth phases. For example, *atpA*, *grpE*, and *tpi* were stable in serum exposure but not during *E. ictaluri* growth. Also, *g6pd* and *mdh* were not stable in serum exposure but were stable during *E. ictaluri* growth. These observations show that stability of a particular reference gene may vary significantly from experiment to experiment, and constant expression levels under different conditions are not warranted and need validation.

In conclusion, selecting suitable reference genes is critical for RT-PCR experiments, and their stable expression should be validated under specific experimental conditions. Interestingly, gene expression studies by RT-PCR and the use of reference genes are not well-known in *E. ictaluri*. This study provides a list of potential reference genes for future gene expression studies in *E. ictaluri* and related bacteria.

